# Cross-reactivity of SARS-CoV structural protein antibodies against SARS-CoV-2

**DOI:** 10.1101/2020.07.30.229377

**Authors:** Timothy A. Bates, Jules B. Weinstein, Scotland E. Farley, Hans C. Leier, William B. Messer, Fikadu G. Tafesse

## Abstract

There is currently a lack of biological tools to study the replication cycle and pathogenesis of SARS-CoV-2, the etiological agent of COVID-19. Repurposing the existing tools, including antibodies of SARS-CoV, is an effective way to accelerate the development of therapeutics for COVID-19. Here, we extensively characterized antibodies of the SARS-CoV structural proteins for their cross-reactivity, experimental utility, and neutralization of SARS-CoV-2. We assessed a total of 10 antibodies (six for Spike, two for Membrane, and one for Nucleocapsid and Envelope viral protein). We evaluated the utility of these antibodies against SARS-CoV-2 in a variety of assays, including immunofluorescence, ELISA, biolayer interferometry, western blots, and micro-neutralization. Remarkably, a high proportion of the antibodies we tested showed cross-reactivity, indicating a potentially generalizable theme of cross-reactivity between SARS-CoV and SARS-CoV-2 antibodies. These antibodies should help facilitate further research into SARS-CoV-2 basic biology. Moreover, our study provides critical information about the propensity of SARS-CoV antibodies to cross-react with SARS-CoV-2 and highlights its relevance in defining the clinical significance of such antibodies to improve testing and guide the development of novel vaccines and therapeutics.

## Introduction

The recent emergence of the novel severe acute respiratory syndrome coronavirus 2 (SARS-CoV-2) in late 2019 has led to an ongoing global COVID-19 pandemic and public health crisis [1]. At the time of writing, there are over seven million confirmed infections and four hundred thousand fatalities globally [2]. SARS-CoV-2 has been designated as a strain of the same species as the original SARS coronavirus (SARS-CoV) due to a high degree of sequence similarity [3]. SARS-CoV-2 falls within the family *Coronaviridae*, and can be further subcategorized as a *Betacoronavirus* of lineage B [3]. There is an urgent need for tools to study this novel coronavirus, as part of the effort to quickly and safely develop vaccines and treatments. One avenue that merits exploration is the repurposing of reagents that were developed for use with SARS-CoV, as many are both extremely effective and commercially available.

Coronaviruses (CoVs) are enveloped, positive-sense, single-stranded RNA viruses with exceptionally large genomes of up to 32 kb on a single RNA molecule. In the case of SARS-CoV-2, two open reading frames code for sixteen nonstructural proteins and other individual open reading frames are responsible for four structural proteins: spike (S), nucleocapsid (N), membrane (M), and envelope (E) and nine accessory proteins [4]. *Coronaviridae* are a large and diverse family of viruses, with several genera further divided into several lineages, and human and animal coronaviruses are intermixed within each of these categories (Forni et al., 2017[5]. Of the human coronaviruses, the SARS-CoVs are most closely related to the lineage C beta-CoV MERS, followed by the lineage A beta-CoVs HCoV-HKU1 and HCoV-OC43, and then the alpha CoVs HCoV-NL63 and HCoV-229E. The lineage A beta-CoVs and the alpha-CoVs are globally distributed with seroprevalence exceeding 90% in some studies, though they cause relatively mild disease compared to the rarer acute respiratory syndrome coronaviruses [6,7].

The four SARS-CoV-2 structural proteins are critical for shaping the physical form of the virion, but most available information about them has been extrapolated from other coronaviruses. Generally, the CoV M protein is involved in shaping the viral envelope membrane [8], the N protein complexes with the viral RNA [9], the S protein mediates receptor recognition and membrane fusion [10,11], and the E protein contributes to the structure of the viral envelope [12]. Furthermore, several of these CoV structural proteins have been shown to have intracellular functions unrelated to their role as structural proteins [9]. There are limits to the utility of extrapolation; it is known, for example, that the topology of the CoV envelope protein varies dramatically between various viruses [12], and the differences between the receptor binding domains (RBDs) of the spike protein can be dramatic. Therefore, tools to interrogate the specific functions of each of the SARS-CoV-2 structural proteins would be of immense and immediate use.

CoV specific antibodies are one type of tool used in such studies. Antibodies against the SARS-CoV-2 structural proteins could be used as reagents in microscopy and western blotting, as structural tools to probe functional epitopes, and even as antiviral therapies. The protein which produces the greatest SARS-CoV-2 specific antibody response in humans is the viral S protein [13], but it is known that antibodies are produced against the N, M, and E proteins as well [7,13]. Since SARS-CoV and SARS-CoV-2 are such markedly similar viruses, as discussed below, it is reasonable to assume that there may be some cross-reactivity between SARS-CoV antibodies against their cognate SARS-CoV-2 structural proteins, and, indeed, there is already some evidence that this is the case [14–17].

SARS-CoV and SARS-CoV-2 S proteins share 76% amino acid sequence homology and both rely on cellular angiotensin-converting enzyme 2 (ACE2) as an attachment receptor as well as the TMPRSS2 protease for priming [18]. Recent reports have identified cross-reactive antibodies that bind to the S protein of both SARS-CoV and SARS-CoV-2, however no such cross-reactive antibodies have been identified for the remaining structural proteins [14–17]. A non-human-primate model of SARS-CoV-2 DNA vaccination found that a polyclonal antibody response to S alone is sufficient to protect from SARS-CoV-2 challenge, similar to results from a human S-only vaccine trial for SARS-CoV [19,20]. Additionally, convalescent plasma from recoverd COVID-19 cases has been broadly shown to reduce mortality of individuals with serious disease [21,22]. The sequence similarity between SARS-CoV and CoV-2 N, M, and E proteins are high, at 91%, 90%, and 95% respectively, making it likely that any individual antibody may be cross-reactive. Indeed, there are reports of human antibodies against the S, N, and M proteins for which the epitopes are identical between SARS-CoV and SARS-CoV-2, further supporting the possibility of cross-reactivity, though none have been experimentally verified [13].

If cross-reactivity with SARS-CoV-2 is a common feature of SARS-CoV antibodies, then many recovered SARS-CoV patients may still possess SARS-CoV-2 reactive antibodies; antibody responses were shown to remain at high levels for at least 12 years according to a recent preprint [23]. While sequence conservation is lower for more common human coronaviruses, their high prevalence may lead to widespread antibodies with cross-reactivity to SARS-CoV-2. Furthermore, antibodies promoting antibody-dependent cellular phagocytosis have been shown to assist in elimination of SARS-CoV infection, showing that cross-reactive antibodies need not be neutralizing to play a productive role in resolution of coronavirus infection [24].

This report characterizes a series of SARS-CoV monoclonal antibodies for cross-reactivity, experimental utility, and neutralization of the SARS-CoV-2 virus. Information about how antibodies from different coronavirus infections interact is critical for several reasons. It is an important factor to consider during the design of antibody-based coronavirus tests, particularly for those as closely related as SARS-CoV and SARS-CoV-2. New treatments for SARS-CoV-2 that interact with a patients immune system will also need to take into account the prevalence of cross-reactive antibodies due to previous coronavirus infections. Further, information about the basic biology of this novel virus will be critical in developing such tailored treatments, and cross-reactive antibodies could be extremely useful in such studies.

## Results

### Sequence similarities of the structural proteins of human coronaviruses

The Biodefense and Emerging Infections (BEI) Research Resources Repository has available several types of antibodies and immune sera against each of the structural SARS-CoV proteins as well as whole virus (summarized in Table 1). Eight of these are mouse monoclonal antibodies of either the IgM, IgG2a, or IgG1 class, recognizing either the SARS-CoV E, M, N, or S proteins. Of these, only two are neutralizing, 341C, and 540C [25]. There are also polyclonal rabbit sera against the SARS-CoV S protein, and an anti-S monoclonal human IgG1 antibody isolated from a SARS-CoV patient [26], all of which are neutralizing.

**Table 1:** SARS-CoV antibodies utilized by this study.

| Antibody | Reference | Protein Specificity | Species | Class | Neutralization of SARS-CoV | Epitope |
| --- | --- | --- | --- | --- | --- | --- |
| 240C | Tripp et al. | S | Mouse | IgG2a | No | S<sub>A</sub> (490-510) |
| 341C | Tripp et al. | S | Mouse | IgG2a | Yes | S<sub>A</sub> (490-510) |
| 540C | Tripp et al. | S | Mouse | IgG2a | Yes | S<sub>A</sub> (490-510) |
| 154C | Tripp et al. | S | Mouse | IgM | No | S<sub>B</sub> (270-350) |
| CR3022 | Ter Meulen et al. | S | Human | IgG1 | Yes | S<sub>C</sub> (369-519) |
| NRC-772 | Made by BEI | S | Rabbit | serum | Yes |  |
| 427C | Tripp et al. | E | Mouse | IgM | No |  |
| 19C | Tripp et al. | M | Mouse | IgM | No |  |
| 283C | Tripp et al. | M | Mouse | IgG1 | No |  |
| 42C | Tripp et al. | N | Mouse | IgM | No |  |

To begin to evaluate structural potential for cross-reactivity, we compared the amino acid sequences of each SARS-CoV-2 structural protein with the homologous protein from the other HCoVs (Figure 1A). We first looked at the amino acid homology among the S proteins of the common human coronaviruses, and found that other human beta-CoVs (MERS-CoV, HCoV-HKU1 and HCoV-OC43) show only about 30% similarity to the SARS-CoV-2 S protein, and human alpha-CoVs (HCoV-229E and HCoV-NL63) show only about 24% similarity to SARS-CoV-2 S protein. The S protein of the original SARS-CoV, however, is much more closely related, showing 77% similarity between SARS-CoV and SARS-CoV-2, which lends support to the idea that anti-SARS-CoV S antibodies could be cross-reactive with the SARS-CoV-2 S protein. The E, M, and N protein sequences show striking similarity between SARS-CoV and SARS-CoV-2; they are 96%, 91%, and 91% similar, respectively (Figure 1A).

**Figure 1:**
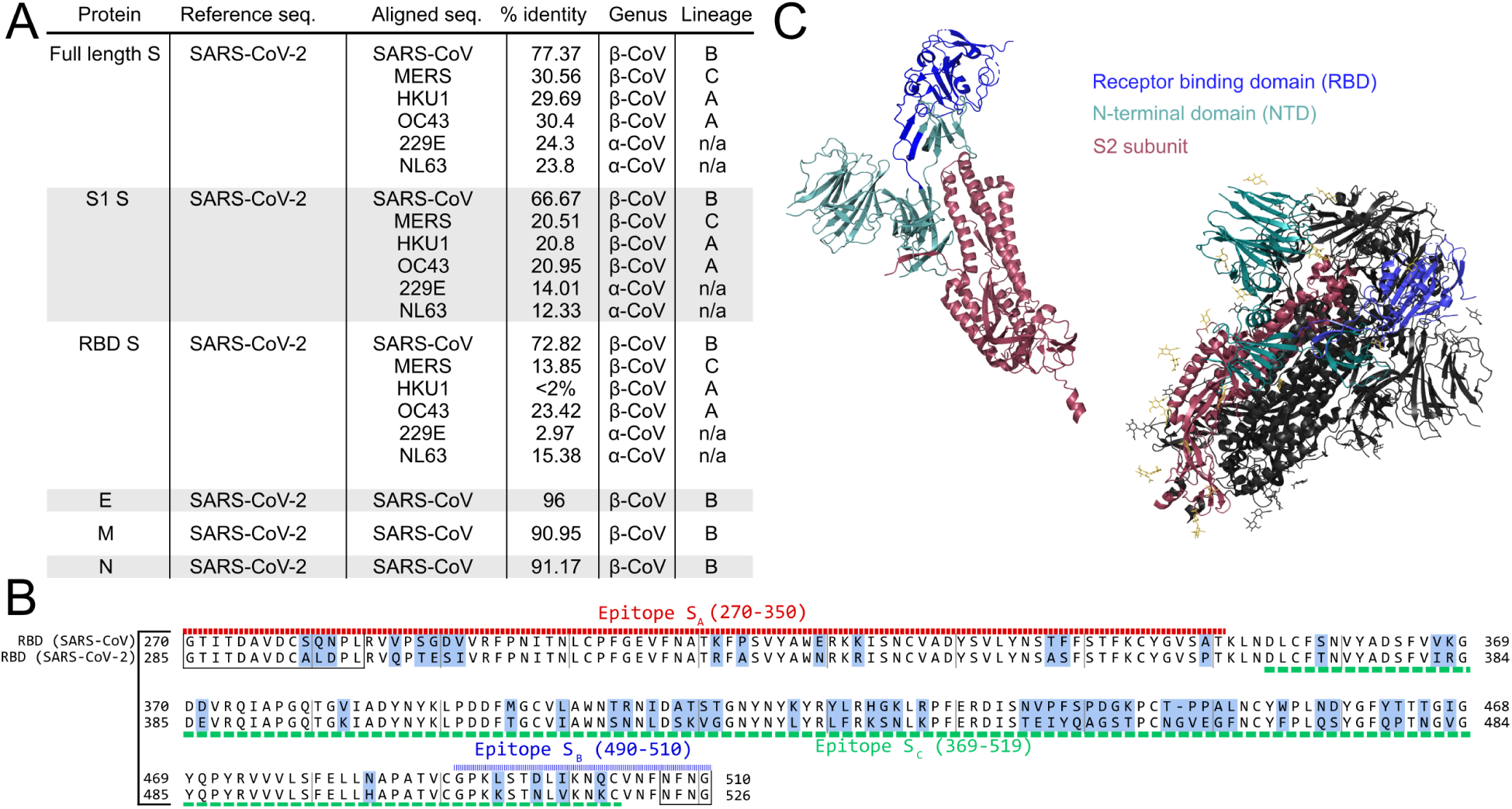
Sequence similarity between SARS-CoV and SARS-CoV-2. (A) Similarity scores for each of the SARS-CoV-2 structural proteins compared with SARS-CoV and other common coronaviruses. Similarity in the S protein is substantially lower than for the other structural proteins. (B) Crystal structure of the S protein color coded by domain both alone and in the functional homotrimeric form where one of the monomers is colored while the other two are in gray. The trimer shows glycosylation sites in yellow. As shown, the NTD and RBD compose the majority of the S1 region. (C) Sequence alignment of the receptor binding domain (RBD) of SARS-CoV and SARS-CoV-2. Regions of difference are highlighted in blue while the epitopes of the antibodies used in this study are underlined according to their designations in Table 1. The boxed regions fall outside of the canonical RBD sequence, but are included due to overlap with the above epitope regions.

While antibodies that recognize each of the structural proteins are of interest as experimental tools, antibodies that recognize the S protein are particularly so because of their potential to neutralize infectious virus. Structural information about the specific biochemical interactions between S-specific antibodies and the S protein is of great value. For the anti-S monoclonal antibodies (through BEI Resources) described in Table 1, the epitopes can be traced to one of three regions of the RBD. While 240C, 341C, and 540C all bind within a region at the end of the RBD (epitope S_A_) [25], the 154C antibody binds to a region at the beginning of the RBD (epitope S_B_); and the human monoclonal antibody CR3022 binds to specific residues in a broad region in the middle of the RBD (epitope S_C_) [17]. These epitopes are indicated in figure 1B, along with the alignment of the SARS-CoV and SARS-CoV-2 RBDs. While not identical, these regions do show some level of similarity between the two virus strains. The three dimensional structure of the spike protein in both monomeric and the functional trimeric form is displayed to illustrate the general accessibility of each portion of the protein (Figure 1C).

### Antibodies of the SARS-CoV structural proteins show cross-reactivities with SARS-CoV-2 by microscopy

To assess SARS-CoV antibodies against SARS-CoV-2, we first performed Immunofluorescence(IF) staining of 293T cells transiently transfected with Strep-tagged constructs of each of the individual SARS-CoV-2 structural proteins [27]. We compared the staining of the Strep-tag within each structural protein in immunofluorescence against that of the experimental antibodies, finding that the staining pattern of a majority of these antibodies as detectable, with some being highly similar to the strep-tag antibody (Figure 2 and S2). The E, M, and S proteins each had at least one high-quality antibody while the only N-specific antibody did not perform well, as was reported for these antibodies against SARS-CoV [25]. Together, these antibodies provide nearly complete coverage of SARS-CoV-2 structural proteins, showing their utility for SARS-CoV-2 experiments involving microscopy.

**Figure 2:**
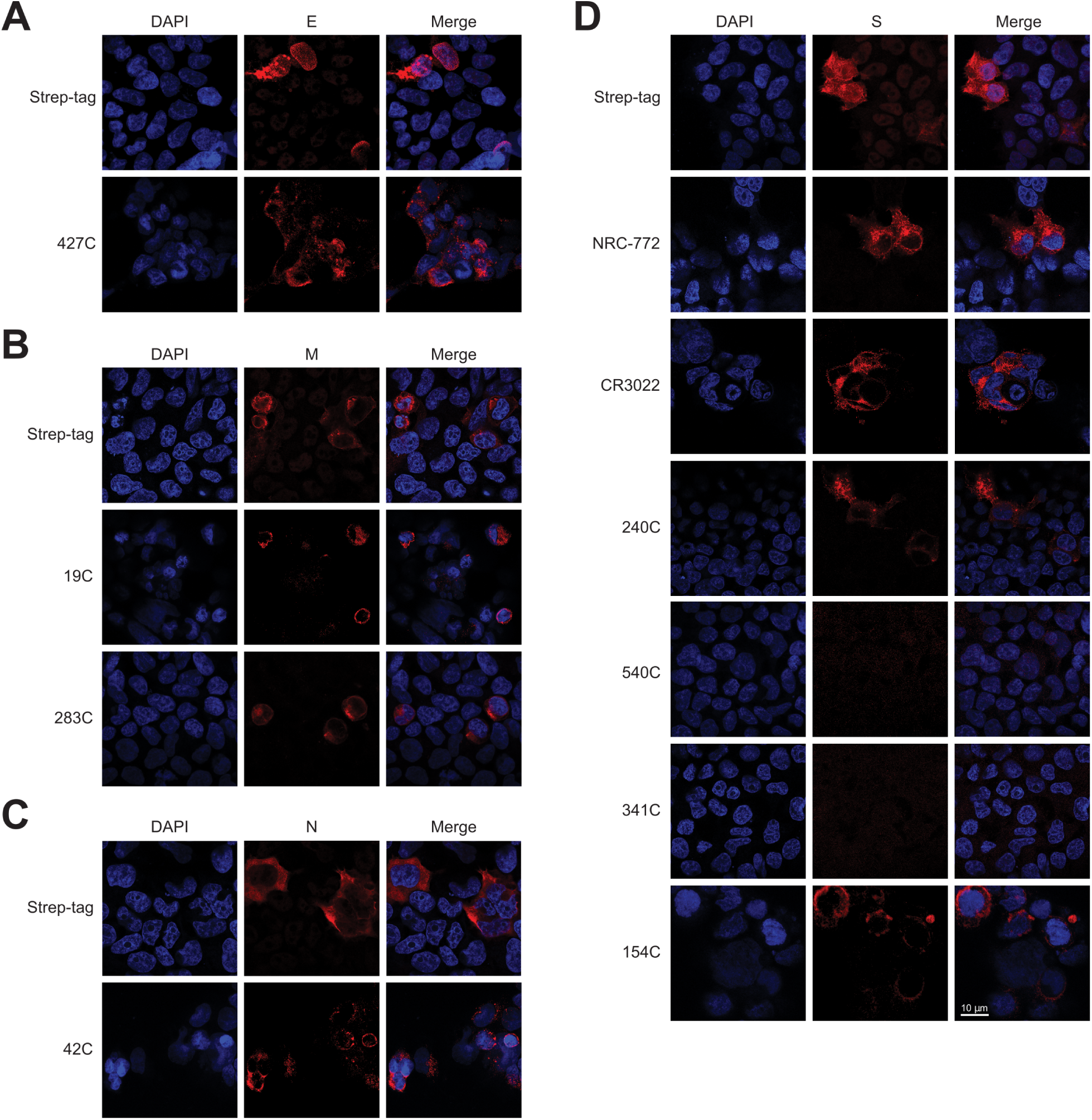
Immunofluorescence of SARS-CoV-2 structural proteins using SARS-CoV antibodies at 63× magnification. Representative immuno-fluorescence images of HEK 293T cells transiently transfected with SARS-CoV-2 structural proteins. 24 h post-transfection, cells were fixed and stained with the listed SARS-CoV antibodies: (A) Envelope, (B) Membrane, (C) Nucleocapsid, and (D) Spike proteins. All proteins are strep-tagged and control stained with anti-strep-tag antibody or the indicated antigen-specific antibody (Red). DAPI (Blue) was used to visualize cell nuclei.

### Antibodies of the SARS-CoV structural proteins show cross-reactivities with SARS-CoV-2 by immunoblotting

We next evaluated these antibodies in western blotting. His_6_-tagged RBD from SARS-CoV-2 was produced in HEK 293 cells and purified by Ni-NTA chromatography (Figure 3A). This RBD was then used for a western blot with each of the mouse monoclonal antibodies (Figure 3B-D). The staining produced by each experimental antibody was compared to anti-His_6_ staining of the RBD as a positive control, and lysate from untransfected 293T cells to assess background. Of these antibodies, 240C and NR-772 produced strong signal with little background, whereas the other monoclonal antibodies did not produce detectable signal (Figure S3).

**Figure 3:**
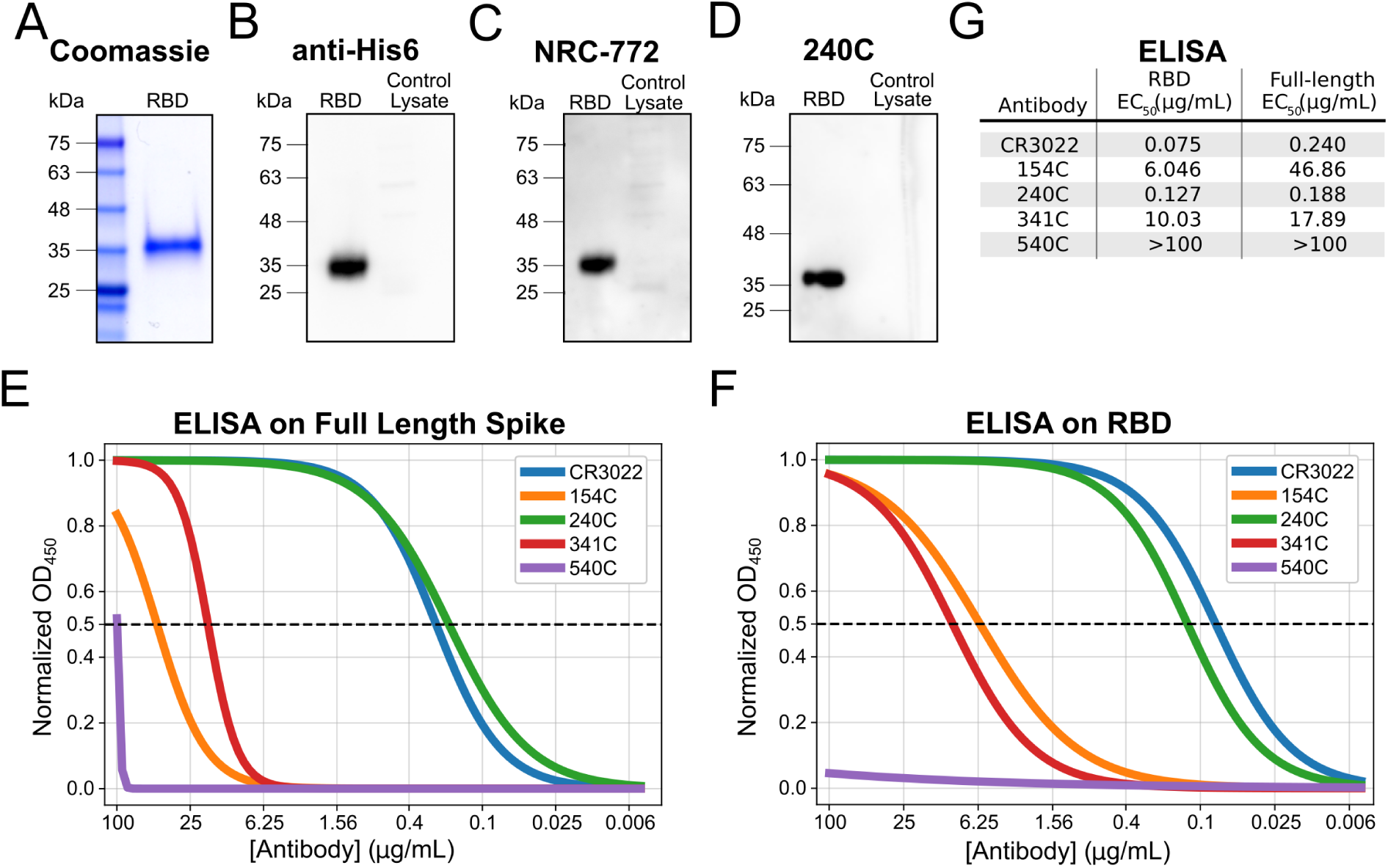
Biochemical characterization of Spike protein specific antibodies. Characterization of the S-specific antibodies by western blot and ELISA. (A) Coomassie stain of in-house purified His6-tagged RBD protein produced in HEK 293-F suspension cells and purified by Ni-NTA chromatography. (B-D) Western blot of purified RBD protein probed with control α-His6 or 240C. 1 μg purified RBD was loaded for each blot and untransfected HEK 293T control lysate was included to monitor non-specific binding. The 240C antibody was the only antibody to demonstrate binding in western blots (see Supplementary Figure 2 for details). (E) ELISA on purified full length Spike coated at 2 μg/mL. (F) ELISA on purified RBD coated at 2 μg/mL. (G) Summary table of observed EC50 values from both sets of ELISAs.

### Spike antibodies show cross-reactivity in binding

The fact that the S glycoprotein is responsible for virus binding and entry into host cells makes it an attractive target for antibody generation as some of these antibodies may be neutralizing. Because of the potential functional role for these antibodies, and because of the number of different antibody clones, we decided to examine the S-protein-specific antibodies more thoroughly.

We assessed the binding of the S-protien-specific antibodies to both the full-length SARS-CoV-2 S protein and the purified Receptor Binding Domain (RBD) by ELISA (Figure 3E-F and Figure S4, summarized in Figure 3G). CR3022 and 240C showed strong binding to both the full-length spike and the RBD (EC_50_ 75 ng/mL and 127 ng/mL, respectively, to the RBD). 154C and 341C showed weak but detectable binding (6.046 µg/mL and 10.03 µg/mL, respectively, to the RBD), while 540C did not demonstrate binding at all. The trend for these antibodies is generally similar to what was seen in the previous studies where the antibodies were tested against recombinant SARS-CoV S protein [25]. The original report of CR3022 did not perform a direct ELISA for us to compare our results to, however our data agree with studies of CR3022 on SARS-CoV-2 showing that it binds strongly to both full-length spike and the RBD [17]. While CR3022 appears to be the strongest binder to RBD, 240C is marginally better on the full-length spike protein.

To assess the binding kinetics of the antibody-RBD interaction in more detail, we measured the antibody-epitope interactions using biolayer interferometry (BLI). The three monoclonal antibodies that showed strongest-binding with the ELISA displayed high affinity for the SARS-CoV-2 RBD, CR3022 showed the strongest binding with a calculated K_D_ of 758 pM (Figure 4A), while 240C demonstrated a 1.36 nM K_D_ (Figure 4B), and 154C a 481 nM K_D_ (Figure 4C). As summarized in figure 4D, these antibodies showed fast-on/slow-off kinetics in agreement with a previous report of CR3022 binding kinetics on RBD [17]. The other antibodies we tested displayed no measurable binding at the highest concentration used (Figure S5). Importantly, BLI does not account for the avidity of these antibodies, and it is likely that the interaction of each epitope/paratope pair is substantially lower than that of the intact antibody; however, the intact antibody more closely resembles the interaction that is likely to occur in most *in vitro* assays, or indeed *in vivo*. Our K_D_ is substantially lower than reported in Tian et al., however this is likely due to differences in the reagents used [15]. Tian et al. expressed their RBD in *E. coli*, preventing glycosylation, while our RBD was produced in mammalian cells. Additionally, their CR3022 was produced as a single chain variable fragment (scFv) in *E. coli*, which would contain only a single paratope and may fold differently than our full CR3022 antibody, which was produced in a plant expression system.

**Figure 1:**
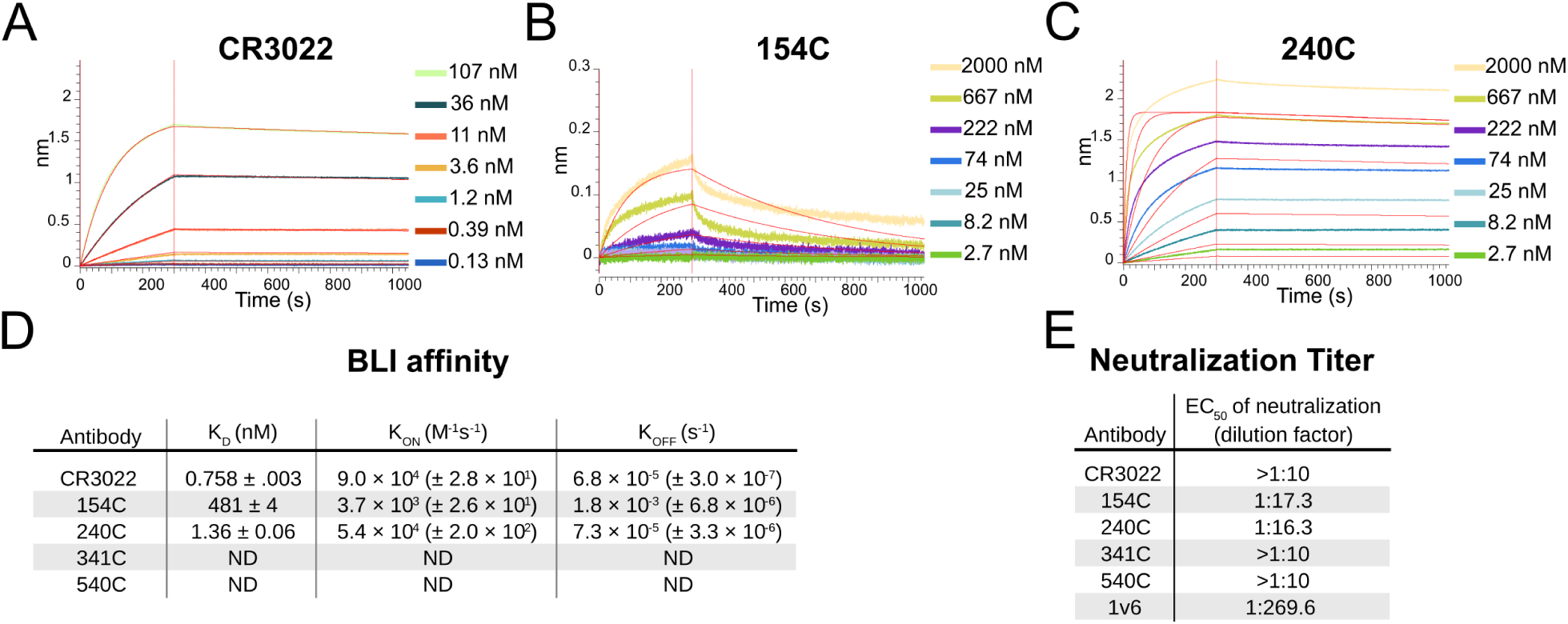
Binding Kinetics and functional testing of Spike specific antibodies against the RBD of the SARS-CoV-2. (A-C) Biolayer interferometry curves for CR3022, 154C, and 240C with three-fold dilutions. Streptavidin biosensors were coated with biotinylated RBD, then blocked with 1 μM D-Biotin in kinetics buffer. Negative binding curves for 341C and 540C shown in figure S4. Curve fitting was performed using 1:1 binding model in ForteBio Analysis HT 10.0 software. (D) Summary of quantified binding kinetics of Spike monoclonal antibodies from BLI experiment. (E) Neutralization assay 50% neutralization values against SARS-CoV-2 Spike pseudotyped lentivirus. 1v6 is positive control from COVID-19 patient convalescent serum collected at day 14. The concentration of all monoclonal antibody stocks was 1 mg/ mL. 154C and 240C showed only partial neutralization at the highest concentration tested (1:10 dilution), while 341C, 540C, and CR3022 failed to reliably neutralize pseudotyped virus at this dilution.

### Spike antibodies of SARS-CoV show limited cross-neutralization of SARS-CoV-2

Finally, we assessed the neutralizing capabilities of these S protein-specific monoclonal antibodies. We set up a neutralization assay using a Lentivirus GFP-reporter pseudotyped with the SARS-CoV-2 S protein [28]. Neutralization was assessed by microscopy, using the area of GFP expression compared to that of an antibody-untreated control. Serial dilutions of antibodies were used to generate neutralization curves and estimate the antibody concentration necessary for 50% neutralization. This readout was used because the monoclonal antibodies displayed only partial neutralization at the highest concentration used in our assay. To validate our assay, we used human convalescent serum from a SARS-CoV-2 positive patient. This anti-serum demonstrated 50% neutralization at a dilution of 1:270 (Figure 4E). Consistent with a previous report, CR3022 failed to show any neutralization at 100 μg/mL despite its potent binding in every other assay [17]. 154C and 240C both showed partial neutralization, with a 50% reduction in GFP area at 57.8 μg/mL and 61.3 μg/mL respectively. Consistent with the BLI results, 341C and 540C did not show substantial neutralization. We were surprised to see 154C perform the best in this assay, particularly because the original report of these antibodies on SARS-CoV showed 341 and 540 as the only antibodies with neutralizing capabilities. One unique aspect of 154C is that it is the only IgM antibody from this selection of Spike-specific antibodies, however it is not clear how this might affect neutralization.

### Summary of cross-reactivity of SARS-CoV structural-protein-specific antibodies to SARS-CoV-2 structural proteins in various assays

The utility of each of the antibodies used in this study has been summarized in Table 2. In particular, the S-protein-specific 240C performed well in every assay we performed, excluding neutralization. In contrast, 540C showed no detectable binding in any of our assays. The other S-protein-specific monoclonal antibodies 154C, 341C, and CR3022 showed mixed utility in different assays. The rabbit polyclonal antibody NRC-772 also worked in every assay in which it was tested, however, polyclonal sera is limited to experiments where structural information about particular epitopes is not important due to the unknown admixture of the contained antibody clones. The antibodies against E, M, and N demonstrated utility in immunofluorescence, but not western blot. Further studies could explore these antibodies in greater detail by producing purified E, M, and N proteins for use in biochemical assays such as the ones we used to characterize the S-protein-specific antibodies in this report.

**Table 2:** Summary of antibody quality for different assays. Qualitative utility scores (+++) very good, (++) good, (+) moderate, (-) fail were assigned to each antibody for each assay in which it was tested; (partial) indicates incomplete neutralization at 100 μg/mL and (ND) indicates that an antibody was not used in this particular assay.

| Antibody | Protein target | Immunofluorescence | ELISA | Western blot | Biolayer interferometry | Neutralization |
| --- | --- | --- | --- | --- | --- | --- |
| 240C | S | +++ | +++ | +++ | +++ | partial |
| 154C | S | + | ++ | - | + | partial |
| 341C | S | + | ++ | - | - | - |
| 540C | S | - | - | - | - | - |
| CR3022 | S | ++ | +++ | - | +++ | - |
| NRC-722 | S | +++ | ND | +++ | +++ | ND |
| 42C | N | - | ND | - | ND | ND |
| 427C | E | ++ | ND | - | ND | ND |
| 19C | M | + | ND | - | ND | ND |
| 283C | M | +++ | ND | - | ND | ND |

## Discussion

Our results demonstrate substantial cross-reactivity from a majority of the SARS-CoV-2 structural-protein-targeted antibodies that we evaluated. This is the first report of cross-reactive antibodies directed against the N, M, and E proteins. These tools can be readily obtained from BEI resources and utilized by labs to study the properties of untagged SARS-CoV-2 structural proteins. These antibodies can serve the unmet need for more resources enabling the study of the SARS-CoV-2. It is critical to understand the basic biology of SARS-CoV-2 in order to inform efforts towards improved diagnostics and treatments. Further, information about cross-reactivity of antibodies between SARS-CoV and SARS-CoV-2 may assist bioinformatics labs in developing computational tools for predicting cross-reactivity of other antibodies, or even guiding rational design of improved coronavirus antibodies and small-molecule therapeutics.

We have shown that these publicly available antibodies are of potential use in several different types of assays with SARS-CoV-2 proteins. We found that several of these SARS-CoV-2 structural protein antibodies demonstrated good staining in immunofluorescence (240C, NRC-772, and CR3022 against S; 283C against M; 427C against E). The anti-S antibodies, 240C and NRC-772, also give clear signal in western blot with minimal background. Several S antibodies show potent binding to full-length S and the RBD by ELISA, as well as binding to the RBD by biolayer interferometry (240C, CR3022, and NRC-772). This wide range of uses substantially broadens our ability to investigate the biochemical properties of SARS-CoV-2 structural proteins.

Although these antibodies only partially neutralized a SARS-CoV-2 model infection, they are still of interest for their potential to elucidate the structure and function of their protein targets. Antibodies have been critical tools in structure determination and in the mapping of proteins’ functional regions. Having a wide array of antibodies that recognize varying epitopes is of great help in this endeavor. Additionally, with the current dearth of knowledge regarding the life cycle and pathogenesis of SARS-CoV-2, particularly regarding the understudied M, N, and E proteins, we believe that these antibodies could be used in experiments to better understand the nuances of their functions beyond their obvious structural roles.

Our results also speak to the high proportion of SARS-CoV antibodies that display substantial cross-reactivity to SARS-CoV-2 structural proteins. Anecdotal evidence supports the efficacy of convalescent plasma treatment for COVID-19, indicating that cross-reactive antibodies generated during previous coronavirus infections may prove beneficial during coronavirus infection [29]. Conversely, studies of COVID-19 patients have found neutralizing antibody titers to be directly proportional to disease severity, suggesting a more complicated relationship between antibodies and COVID-19 [30,31]. Some have hypothesized that this may be due to high concentrations of virus and neutralizing antibodies acting together to drive greater immune pathology [32,33]. A better understanding of the functions of individual antibody isotypes against different antigenic targets will be critical to predicting the utility of a particular antibody against SARS-CoV-2.

Further studies could also investigate possible cooperation between antibodies recognizing different epitopes, especially as CR3022 neutralization was shown to have synergy with another anti-S antibody which recognized a different epitope on the protein [26]. A recent study [34], for example, characterized a neutralizing monoclonal antibody that did not bind the RBD at all, and instead recognized an epitope in the NTD of the S protein. Knowledge about the variety of vulnerable epitopes, and possible synergy between antibodies that target them, brings us ever closer to being able to design and deploy effective therapeutics and vaccines in this time of urgent need.

## STAR★ Methods

| Reagent or Resource | Identifiers | Source |
| --- | --- | --- |
| **Antibody** |  |  |
| Human monoclonal anti-SARS-CoV -S Cr3022 | NR-52392 | BEI Resources |
| Mouse anti-SARS-CoV S Monoclonal IgM 154c | NR-620 | BEI Resources |
| Mouse anti-SARS-CoV S Monoclonal IgG2a 240c | NR-616 | BEI Resources |
| Mouse anti-SARS-CoV S Monoclonal IgG2a 341c | NR-617 | BEI Resources |
| Mouse anti-SARS-CoV S Monoclonal IgG2a 540c | NR-618 | BEI Resources |
| Mouse anti-SARS-CoV M Monoclonal IgM 19c | NR-615 | BEI Resources |
| Mouse anti-SARS-CoV M Monoclonal IgG1 283c | NR-621 | BEI Resources |
| Mouse anti-SARS-CoV E Monoclonal IgM 472c | NR-614 | BEI Resources |
| Mouse anti-SARS-CoV N Monoclonal IgM 42c | NR-619 | BEI Resources |
| Rabbit anti-SARS-CoV S polyclonal sera-hydrogel | NRC-773 | BEI Resources |
| Rabbit anti-SARS-CoV S polyclonal sera | NRC-768 | BEI Resources |
| Guinea pig anti-SARS-CoV S polyclonal sera | NR-10316 | BEI Resources |
| Mouse anti-2xStrep monoclonal | SAB2702215 | Sigma Aldrich |
| antibody |  |  |
| Goat anti-mouse HRP IgG | HAF007 | R & D Systems |
| Goat anti-mouse HRP IgM | 62-6820 | Invitrogen |
| Donkey anti-human HRP | SAB3701359 | Sigma Aldrich |
| antiHis-HRP | ma1-80218 | Invitrogen |
| anti-Human-AlexaFluor488 | A11013 | Invitrogen |
| anti-Mouse IgG AlexaFluor555 | A21422 | Invitrogen |
| anti-Mouse IgM AlexaFluor488 | A21042 | Invitrogen |
| anti-Rabbit FAB2 AlexaFluor555 | 4413S | Invitrogen |
| **Biological samples** |  |  |
| Human deidentified Patient Sera 1v6 | N/A | Messer Lab, OHSU MMI |
| Human deidentified Patient Sera 0v1 | N/A | Messer Lab, OHSU MMI |
| **Chemicals and recombinant proteins** |  |  |
| RBD recombinant purified Sars-CoV-2 Spike | NR-52306 | BEI Resources |
| Full-length Soluble SARS-CoV-2 Spike | NR-52308 | BEI Resources |
| BSA |  | Gold-Bio |
| FBS | 97068-085 | VWR |
| PBS | 17-516F | Lonza Biosciences |
| Trypsin | 25200056 | Thermo Fisher Scientific |
| Recombinant spike protein (His tag),SARS-CoV-2 | NR-52309 | BEI Resources |
| **Cell line** |  |  |
| HEK 293T | CRL-3216 | ATCC |
| HEK 293T human Ace2 | NR-52511 | Crawford et al. 2020 , |
| expressing stable cell line |  | Bloom lab, BEI Resources |
| **Recombinant DNA** |  |  |
| HDM_IDTSpike_fixK, SARS-CoV-2 plasmid | NR-52514 | Crawford et al. 2020 ,<br>Bloom lab, BEI Resources |
| HDM_Hgpm2 | NR-52517 | Crawford et al. 2020 ,<br>Bloom lab, BEI Resources |
| HDM_tat1b | NR-52518 | Crawford et al. 2020 ,<br>Bloom lab, BEI Resources |
| pRC_CMV_Rev1b | NR-52519 | Crawford et al. 2020 ,<br>Bloom lab, BEI Resources |
| pHAGE2_CMV_ZsGreen_W | NR-52520 | Crawford et al. 2020 ,<br>Bloom lab, BEI Resources |
| pTwist-EF1alpha-nCoV-2019-S-2xStrep |  | Gordon et al. 2020<br>Krogon Lab |
| pLVX-EF1alpha-nCoV-2019-E-IRES-Puro |  | Gordon et al. 2020<br>Krogon Lab |
| pLVX-EF1alpha-nCoV-2019-M-IRES-Puro |  | Gordon et al. 2020<br>Krogon Lab |
| pLVX-EF1alpha-nCoV-2019-N-IRES-Puro |  | Gordon et al. 2020<br>Krogon Lab |
| Plasmid pCAGGS, Spike protein(soluble),SARS-CoV-2,w/his tag | NR-52309 | BEI Resources |
| Plasmid pcDNA3.1 SARS-CoV-2 Spike ectodomain (His) | NR-52421 | BEI Resources |
| Plasmid pMCSG53 SARS-CoV-2, Spike RBD (His) | NR-52430 | BEI Resources |

| Software |  |  |
| --- | --- | --- |
| R | N/A | rstudio.com |
| Python | N/A | python.org |
| Critical Commercial Assays |  |  |
| PureLink™ HiPure Plasmid<br>Maxiprep Kit | K210007 | Invitrogen |
| ChromaLINK biotin protein<br>labeling kit | B-9007-105K | Vector Labs |
| Streptavidin (SA) Biosensors | 18-5019 | ForteBio |
| Other |  |  |
| NuncSorp ELISA Plates |  | Thermo Fisher Scientific |
| Nitrocellulose membrane |  |  |

## Resource Availability

### Lead contact

Further information and requests for resources and reagents should be directed to and will be fulfilled by the Lead Contact, Fikadu Tafesse.

### Materials availability

No unique reagents were generated during the course of this study.

### Data and software availability

This study did not generate any unique datasets or code.

## Experimental Model and Subject Details

293T stable cell lines expressing Ace2 receptor (293T-Ace2) were a kind gift from Dr. Jesse D. Bloom from University of Washington, and described previously (Crawford et al. 2020). Wt low-passage 293T cells (293T-Lp) and 293T-Ace2 were cultured in Dulbecco’s Modified Eagle Medium (DMEM, 10% FBS, 1% Penn-Strep, 1% NEAA) at 37C. Cells were cultured on treated T75 dishes, passaged with Trypsin at 95% confluency to avoid overcrowding.

## Method Details

### Sequence alignment

Protein sequences were obtained from uniprot and aligned using the T-Coffee multiple sequence alignment server.

### Cell transfection

Transfections were carried out in 293T cells seeded at 70-90% cell density using Lipofectamine 3000 (ThermoFisher Scientific) as per manufacturer’s instructions. For immunofluorescence, the SARS-CoV2 structural protein plasmids pTwist-EF1alpha-nCoV-2019-S-2xStrep, pLVX-EF1alpha-nCoV-2019-E-IRES-Puro, pLVX-EF1alpha-nCoV-2019-M-IRES-Puro, or pLVX-EF1alpha-nCoV-2019-N-IRES-Puro were transfected using 2μg of plasmid per well of a 24-well plate. Structural SARS-CoV-2 protein plasmids were a kind gift from the Krogan Lab at UCSF and are described previously (Gordon et al 2020). For pseudotyped lentivirus production, lentivirus packaging plasmids, HDM_Hgpm2, HDM_tat1b, PRC_CMV_Rev1b, SARS_CoV-2 S plasmid HDM_IDTSpike_fixK, and LzGreen reporter plasmid pHAGE2_CMV_ZsGreen_W were transfected using 0.44μg for packaging, 0.68μg for S, and 2μg for reporter plasmids per 6 cm dish. Packaging, SARS-CoV-2 S, and reporter plasmids were a kind gift from Jesse D. Bloom from University of Washington, and are described previously (Crawford et al 2020). Transfection media was carefully removed 6 hours post transfection, and replaced with DMEM.

### Pseudotyped lentivirus production

293T cells were seeded at 2 million cells/dish in 6cm TC-treated dishes. The following day, cells were transfected as described above with lentivirus packaging plasmids, SARS-CoV-2 S plasmid, and lzGreen reporter plasmid (Crawford et al., 2020). After transfection, cells were incubated at 37C for 60 hours. Viral media was harvested, filtered with 0.45μm filter, then frozen before use. Virus transduction capability was then tittered on 293T-Ace2 cells treated with 50μl of 5μg/ml polybrene (Sigma-Aldritch LLC). LzGreen titer was determined by fluorescence using BZ-X700 all-in-one fluorescent microscope (Keyence), a 1:16 dilution was decided as optimal for following neutralization assays due to broad transduced foci distribution.

### Immunofluorescence

293T cells were seeded on 24-well plates containing glass coverslips coated with poly-lysine solution; 100,000 cells were seeded per well. Cells were transfected with SARS-CoV-2 structural protein plasmids as described above. After 48 hours post transfection, cells were fixed with 4% PFA in PBS. Cover slips were permeabilized with 2% BSA, 0.1% Triton-X-100 in PBS. Transfected cells were incubated for 3 hours at RT with the following anti-SARS-CoV structural protein monoclonal or polyclonal antibodies at a 1:250 dilution: mouse anti-SARS-CoV S monoclonal IgM 154c, mouse anti-SARS-CoV S monoclonal IgG2a 240c, mouse anti-SARS-CoV S monoclonal IgG2a 341c, mouse anti-SARS-CoV S monoclonal IgG2a 540c, mouse anti-SARS-CoV N monoclonal IgM 19c, mouse anti-SARS-CoV M monoclonal IgG1 283c, mouse anti-SARS-CoV E monoclonal IgM 472c, mouse anti-SARS-CoV N monoclonal IgN 42c, rabbit anti-SARS-CoV S polyclonal sera (BEI Resources) and mouse anti-2xStrep-tag antibody (Sigma-Aldrich). Anti-mouse IgG AF555, anti-rabbit IgG AF555, or anti-mouse IgM AF488 conjugated secondary antibodies were added at 1:500 dilution for 1 hour at RT (Invitrogen). Confocal imaging was performed with a Zeiss LSM 980 using a 63x Plan-Achromatic 1.4 NA oil immersion objective. Images were processed with Zeiss Zen Blue software. Maximum intensity *z*-projections were prepared in Fiji. All antibody stain images were pseudocolored red for visual consistency.

### Neutralization assay

Neutralization protocol was based on previously reported neutralization research utilizing SARS-CoV-2 S pseudotyped lentivirus (Crawford et al., 2020). 293T-Ace2 cells were seeded on tissue culture treated, poly-lysine treated 96-well plates at a density of 10,000 cells per well. Cells were allowed to grow overnight at 37°C. LzGreen SARS-COV-2 S pseudotyped lentivirus were mixed with 2-fold dilutions of the following monoclonal or polyclonal anti-SARS-CoV-2 S antibodies: mouse anti-SARS-CoV S monoclonal IgM 154c, mouse anti-SARS-CoV S monoclonal IgG2a 240c, mouse anti-SARS-CoV S monoclonal IgG2a 341c, mouse anti-SARS-CoV S monoclonal IgG2a 540c, rabbit anti-SARS-CoV S polyclonal sera, Guinea pig anti-SARS-CoV S polyclonal sera, human monoclonal anti-SARS-CoV S Cr3022 (BEI Resources). Human patient sera from a SARS-CoV-2 patient was used as positive neutralization control, while virus alone was used as negative control. Sera and antibody dilutions ranged from 1:10 to 1:1048. Virus-antibody mixture was incubated at 37C for 1 hour after which virus was added to 293T-Ace2 treated with 5μg/ml polybrene. Cells were incubated with neutralized virus for 44 hours before imaging. Cells were fixed with 4% PFA for 1 hour at RT, incubated with DAPI for 10 minutes at RT, and imaged with BZ-X700 all-in-one fluorescent microscope (Keyence). Estimated area of DAPI and GFP fluorescent pixels was calculated with built in BZ-X software (Keyence).

### Enzyme-linked immunosorbent assay (ELISA)

ELISA plates, Nunc MaxiSorp (Invitrogen), were coated with purified recombinant SARS-COV2 RBD domain (BIR resources, NR-52306) at 2μg/ul in PBS. Coating was carried out overnight at 4°C. Protein was blocked in 2% BSA, 1% tween-20 in PBS for 30 minutes at RT. The following anti SARS-CoV-2 S monoclonal and polyclonal antibodies were serially diluted by 2-fold dilutions in blocking buffer: mouse anti-SARS-CoV S monoclonal IgM 154c, mouse anti-SARS-CoV S monoclonal IgG2a 240c, mouse anti-SARS-CoV S monoclonal IgG2a 341c, mouse anti-SARS-CoV S monoclonal IgG2a 540c, human monoclonal anti-SARS-CoV-S Cr3022 (BEI Resources). Human patient sera from a SARS-CoV-2 patient was used as a positive control. Dilutions ranged from1:10 to 1:10480, and were incubated for 1 hour at RT. Anti-mouse HRP, and anti-human-HRP secondary antibodies were used at 1:4000 concentration in blocking buffer, and were incubated 1 hour at RT. 50 μL of TMB HRP substrate (ThermoFisher Scientific) was added, and following incubation for 10 minutes at RT, 50μL of 2N H2SO4 was added as a stopping solution. Plate absorbance at 405nm was measured using a CLARIOstar® Plus plate fluorimeter (BMG Labtech).

### RBD protein purification and biotinylation

Purified SARS-CoV-2 S-RDB protein was prepared as described previously (Stadlbauer et al., 2020). Briefly, His-tagged RBD bearing lentivirus was produced in HEK 293T cells and used to infect HEK 293-F suspension cells. The suspension cells were allowed to grow for 3 days with shaking at 37°C at 8% CO_2_. Cell supernatant was collected, sterile filtered, and purified by Ni-NTA chromatography. The purified protein was then buffer exchanged into PBS and concentrated. For use in BLI, purified RBD was biotinylated using the ChromaLINK biotin protein labeling kit according to the manufacturer’s instructions with 5x molar equivalents of labeling reagent to achieve 1.92 biotins/protein.

### Biolayer interferometry (BLI)

Streptavidin biosensors (ForteBio) were soaked in PBS for at least 30 minutes prior to starting the experiment. Biosensors were prepared with the following steps: equilibration in kinetics buffer (10 mM HEPES, 150 mM NaCl, 3mM EDTA, 0.005% Tween-20, 0.1% BSA, pH 7.5) for 300 seconds, loading of biotinylated RBD protein (10ug/mL) in kinetics buffer for 200 seconds, and blocking in 1 μM D-Biotin in kinetics buffer for 50 seconds. Binding was measured for seven 3-fold serial dilutions of each monoclonal antibody using the following cycle sequence: baseline for 300 seconds in kinetics buffer, association for 300 seconds with antibody diluted in kinetics buffer, dissociation for 750 seconds in kinetics buffer, and regeneration by 3 cycles of 20 seconds in 10 mM glycine pH 1.7, then 20 seconds in kinetics buffer. All antibodies were run against an isotype control antibody at the same concentration. Data analysis was performed using the ForteBio data analysis HT 10.0 software. Curves were reference subtracted using the isotype control and each cycle was aligned according to its baseline step. KDs were calculated using a 1:1 binding model using global fitting of association and dissociation of all antibody concentrations, excluding dilutions with response below 0.005 nm.

### Western blot

293T cells were seeded in 10 cm dishes at a density of 3.5 million cells per dish. After overnight growth, cells were transfected using lipofectamine 3000 as described above. Plasmids pTwist-EF1alpha-nCoV-2019-S-2xStrep, pLVX-EF1alpha-nCoV-2019-E-IRES-Puro, pLVX-EF1alpha-nCoV-2019-M-IRES-Puro, or pLVX-EF1alpha-nCoV-2019-N-IRES-Puro were transfected using 90 μg of DNA per 10 cm dish. Cells were scraped 48 hours post-transfection, then lysed in RIPA buffer (EMD Millipore). Cell lysates were diluted with reducing Laemmli buffer, incubated for 10 minutes at 37°C, then ran on 4– 20% Mini-PROTEAN^®^ TGX™ Precast Protein Gels (BIO-RAD). Additionally, 1ug of purified recombinant S RBD-His_6_ was diluted in PBS and Laemmli buffer to a final volume of 20 μl and added to a 7.5% Mini-PROTEAN^®^ TGX™ Precast Protein Gel (BIO-RAD). Resolved proteins were then transferred to a PVDF membrane, blocked in TBS with 2% BSA 0.1% Tween-20, then incubated with the following antibodies diluted to 1:500 in blocking buffer: mouse anti-SARS-CoV N monoclonal IgM 19c, mouse anti-SARS-CoV M monoclonal IgG1 283c, mouse anti-SARS-CoV E monoclonal IgM 472c, and mouse anti-2xStrep-tag antibody, and anti-His-HRP. Blots were stained with SuperSignal™ West Pico PLUS Chemiluminescent Substrate (ThermoFisher Scientific) using an ImageQuant LAS 4000 imager (GE Life Sciences).

## Quantification and Statistical Analysis

The EC_50_ values were calculated using a three parameter logistic regression model in the Python software package.

## Acknowledgments

This work was supported by NIH training grant T32AI747225 on Interactions at the Microbe-Host Interface and OHSU Innovative IDEA grant 1018784. BLI data was generated on an Octet Red 384, which is made available and supported by the OHSU Proteomics Shared Resource facility and equipment grant number S10OD023413. We acknowledge the support of the members of the Messer lab who performed collection of patient samples, and the patients who agreed to donate samples for scientific research.

## Author Contributions

T.A.B., J.W., and F.G.T. conceived and designed the study. T.A.B., J.W., and H.L. performed the experiments and collected the data. T.A.B. performed data analysis and visualization. T.A.B., S.F., and J.W. wrote the paper, and all authors reviewed and edited the paper.

