## Supplemental Figures for "Cross-reactivity of SARS-CoV structural protein antibodies against SARS-CoV-2"

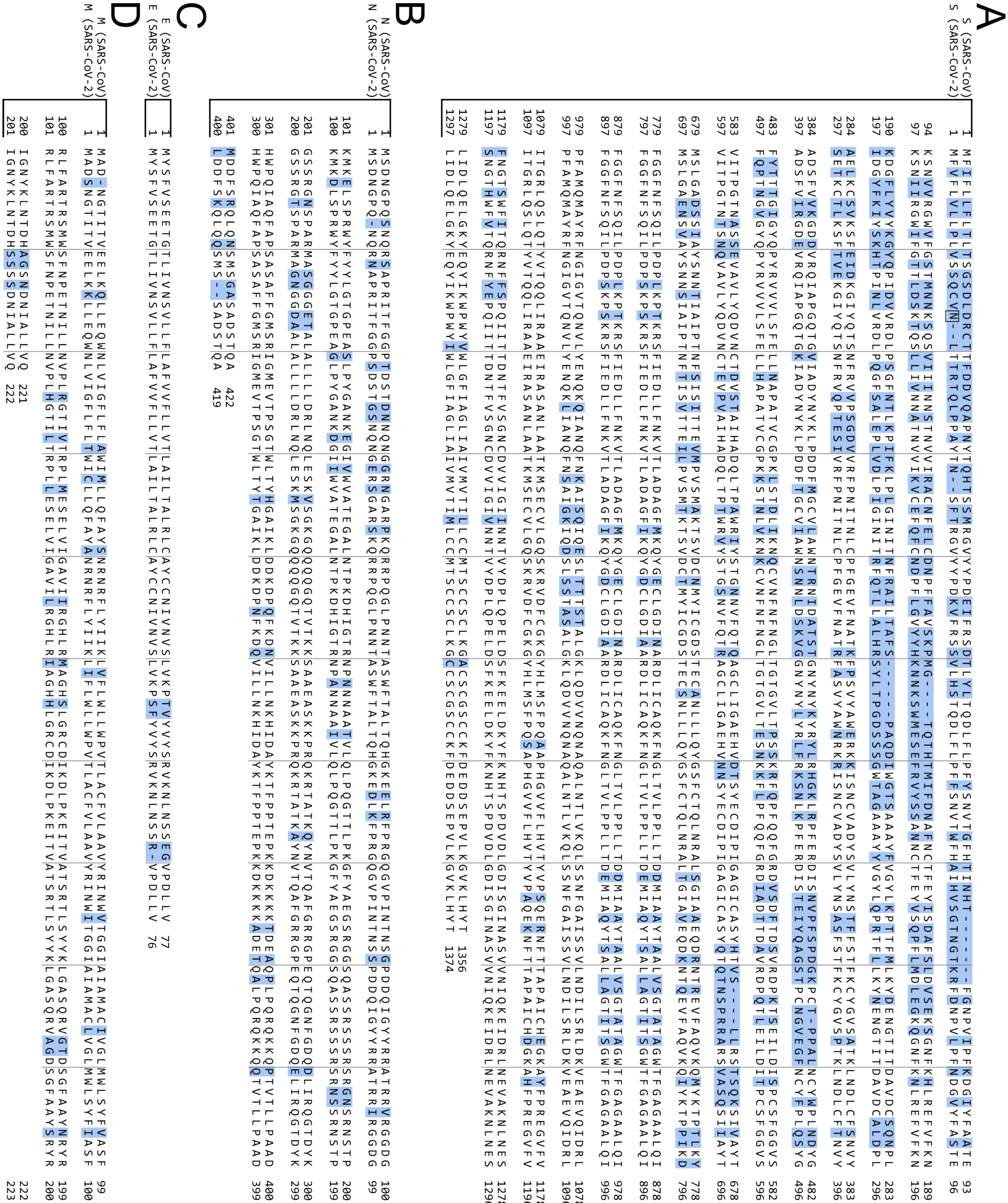


Figure S1: Alignment of structural proteins for SARS-CoV and SARS-CoV-2. Protein sequences were obtained from Uniprot. Differences are highlighted in blue. Grey lines are spaced every 10 characters. For each pair, SARS-Cov is on the top and SARS-CoV-2 is on the bottom.

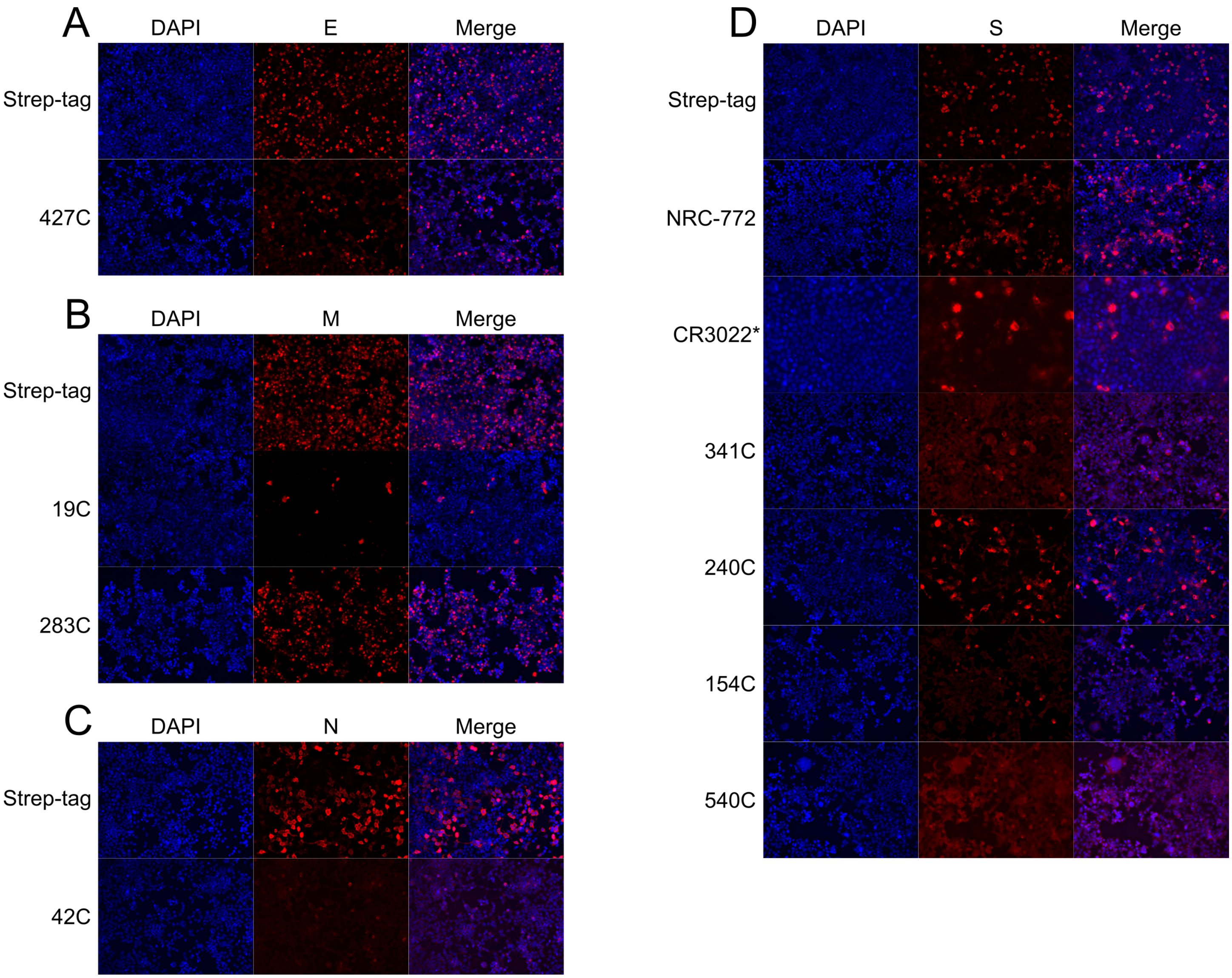


Figure S2: Immunofluorescence of SARS-CoV-2 structural proteins using SARS-CoV antibodies at 10× magnification. Representative immuno-fluorescence images of HEK 293T cells transiently transfected with SARS-CoV-2 structural proteins. 24 h post-transfection, cells were fixed and stained with the listed SARS-CoV antibodies: (A) Envelope, (B) Membrane, (C) Nucleocapsid, and (D) Spike proteins. All proteins are strep-tagged and control stained with anti-strep-tag antibody or the indicated antigen-specific antibody (Red). DAPI (Blue) was used to visualize cell nuclei. * indicates that CR3022 was imaged at 20× magnification.

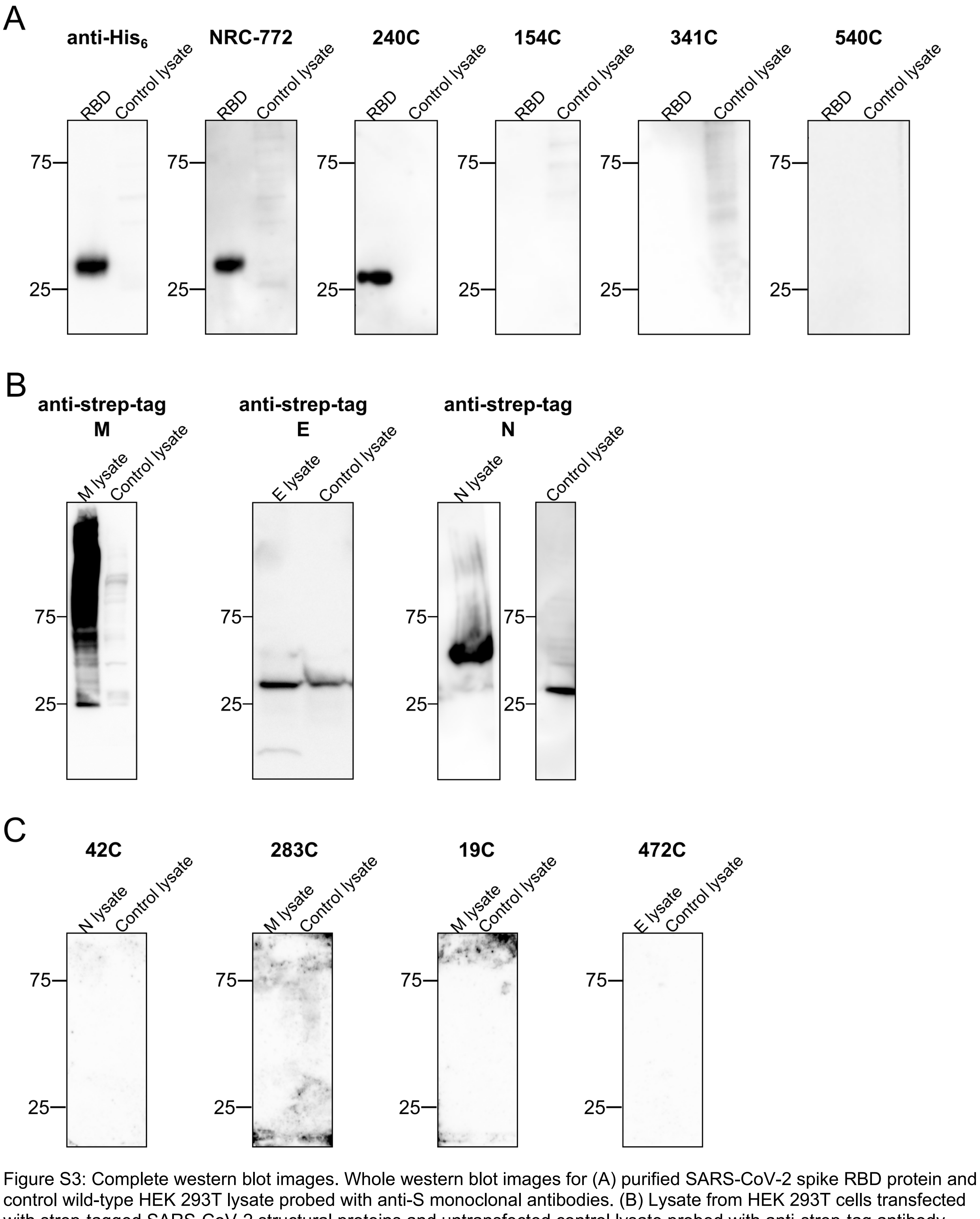


Figure S3: Complete western blot images. Whole western blot images for (A) purified SARS-CoV-2 spike RBD protein and control wild-type HEK 293T lysate probed with anti-S monoclonal antibodies. (B) Lysate from HEK 293T cells transfected with strep-tagged SARS-CoV-2 structural proteins and untransfected control lysate probed with anti-strep-tag antibody. (C) Transfected and control lysate western blots probed with monoclonal antibodies specific to N, M, or E.

A

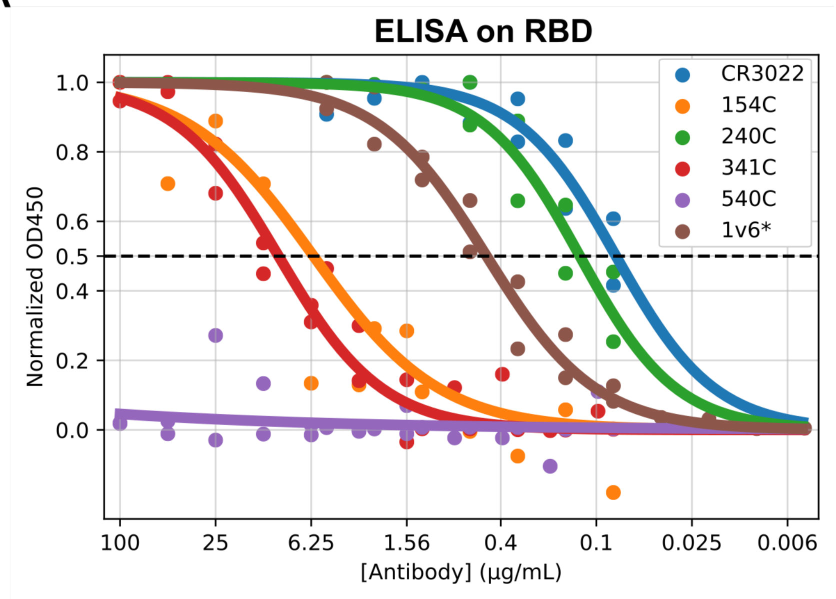


B

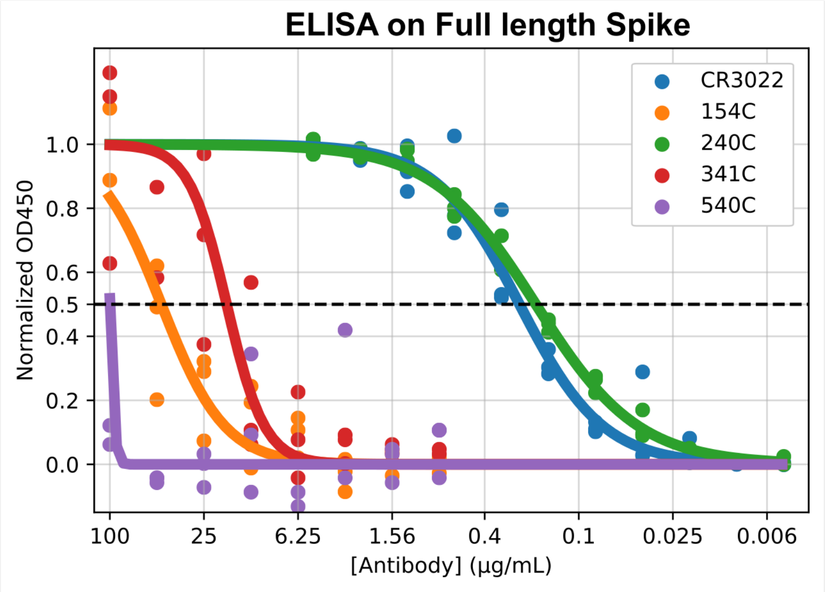


Figure S4: ELISA extended data. ELISA against (A) RBD coated at 2 µg/mL and (B) full length spike coated at 2 µg/mL and then probed with the indicated monoclonal antibody, or 1v6 human convalescent serum. Each point represents the mean of 2 or 3 technical replicates from a single experiment. Data were normalized according to the maximum signal seen for each secondary antibody in each experiment. *1v6 is convalescent serum used to validate the assay. A stock concentration of 1 mg/mL was used to facilitate calculations for reference purposes, but this does not represent an accurate $EC_{50}$ value.

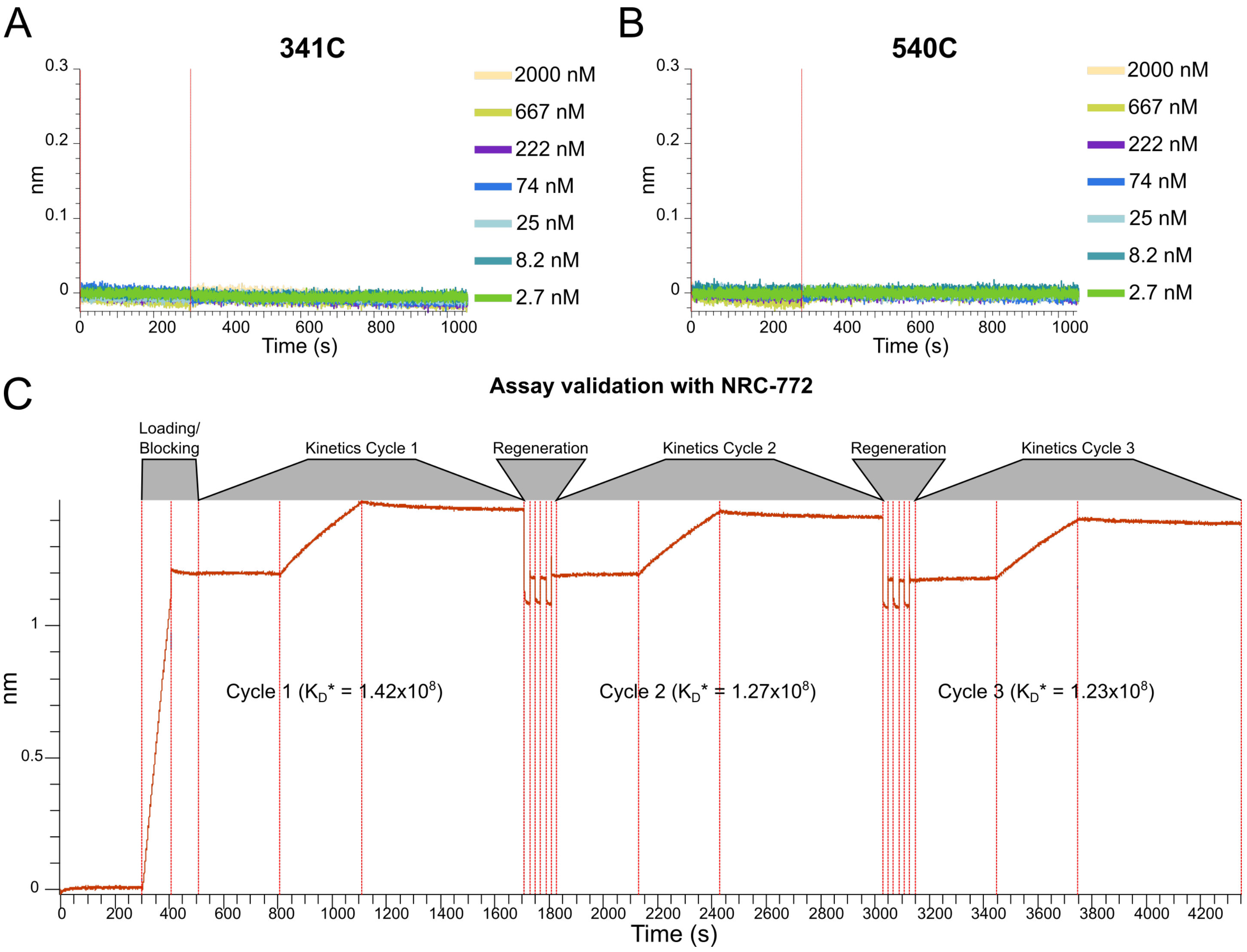


Figure S5: BLI extended data. Negative binding curves for antibodies (A) 341C and (B) 540C. (C) NRC-772 rabbit polyclonal antibody used for method validation. Curves show minimal loss of signal with multiple regeneration cycles as well as stable $K_D$ values, demonstrating stability of RBD under regeneration conditions. NRC-772 serum used at 1:50 dilution in kinetics buffer. *$K_D$ values assume 1 mg/mL initial concentration in order to facilitate $K_D$ calculation only as a reference between cycles and does not represent an accurate affinity measurement.
